# Stacking of PRRs in potato to achieve enhanced resistance against *Phytophthora infestans*

**DOI:** 10.1101/2023.09.07.556738

**Authors:** Yerisf C. Torres Ascurra, Doret Wouters, Richard G. F. Visser, Thorsten Nürnberger, Vivianne G. A. A. Vleeshouwers

## Abstract

Plants employ pattern recognition receptors (PRRs) to sense pathogen-associated molecular patterns (PAMPs) or apoplastic effectors at the plant cell surface, as well as nucleotide-binding domain leucine-rich-repeat-containing receptors (NLRs) to sense effectors inside the plant cell. Breeding for potato resistance to *P. infestans* has focused on the use of NLRs, however, these genes are typically quickly overcome since the matching avirulence genes evolve exceptionally quickly. Here, we stacked two PRRs, *PERU* and *RLP23*, that recognize the rather conserved *Phytophthora* PAMPs Pep-13/25 and nlp20, respectively, in the potato cultivar Atlantic, and evaluated their effect on *P. infestans* resistance. We found that PERU and RLP23 cooperate for the early immune responses like the accumulation of reactive oxygen species (ROS) and production of ethylene by recognizing their corresponding PAMPs. Furthermore, we show that potato plants overexpressing these two PRRs are slightly less affected by *P. infestans* compared to the single transformants. Together, our data suggest that pyramiding of surface receptors can provide additional enhanced resistance against pathogens, however, more effective or synergistic combinations that may include intracellular NLR receptors should be explored.

---

Potato (*Solanum tuberosum* L.) is one of the most important food crops worldwide with a production of more than 359 million tons in 2020 [1]. However, potato is severely affected by the late blight disease with yield losses estimated at 15-20% globally [2]. *Phytophthora infestans*, the causal agent of late blight, is a very destructive oomycete that can dismantle a complete potato field within days [3]. *P. infestans* has remained the most important threat for potato production, mainly because of its elevated virulence and distinctive genome architecture that underpins its evolutionary potential and ability to rapidly adapt to resistant plants [4, 5].

Breeding for resistance against *P. infestans* in potato has been focused on the use of intracellular nucleotide-binding domain leucine-rich-repeat-containing receptors (NLRs) encoded by resistance (*R*) genes. NLRs activate immune responses upon recognition of effector proteins secreted by pathogens, leading to effector-triggered immunity (ETI) [6]. Almost 50 *R* genes have been identified from wild tuber-bearing relatives of potato [7]; however, most, if not all, *R* genes have been overcome by *P. infestans* because of the rapid evolution of the matching avirulence (Avr) effectors, by diverse mechanisms such as mutation, gene loss, suppression and gene silencing that leads to loss of recognition [4]. Some recently identified genes including *RB/Rpi-blb1, Rpi-vnt1, Rpi-Smira2*/*R8, Rpi-blb2* confer broad-spectrum resistance to most of the tested *P. infestans* strains, according to long term field assays [8, 9]. Also *Rpi-amr1* from *Solanum americanum* was found to confer resistance to all 19 tested *P. infestans* isolates in transgenic potato [10]. However, the durability of these newer *Rpi* genes remains to be evaluated after prolonged deployment at larger acreages.

Despite the quick evolution of virulent races, stacking of *R* genes is considered an effective strategy to obtain a more durable resistance against *P. infestans* in potato. Three resistance genes to *P. infestans* (*Rpi*), *Rpi-sto1, Rpi-vnt1*.*1* and *Rpi-blb3* from the potato wild relatives *S. stoloniferum, S. venturii* and *S. bulbocastanum*, respectively, were placed into a single binary vector PBINPLUS and transformed to the cultivar Désirée, which led to a broad resistance spectrum, with no silencing effects observed under laboratory conditions [11]. These transgenic plants (and plants with single *R* gene) were evaluated in field trials in The Netherlands and Belgium for two years. Only the plants with the stacked genes remained healthy until the end of the season, while plants with single *R* genes showed disease symptoms [12]. In another study, the *NLR* genes *Rpi-blb1, Rpi-blb2* from *S. bulbocastanum* and *Rpi-vnt1*.*1* from *S. venturii* were cloned in one transcriptional unit and transformed into the cultivars Désirée and Victoria. The transgenic events showed complete resistance to *P. infestans* in field trials in Uganda during three consecutive seasons [13]. Still, the durability of the resistance obtained by gene stacking should be carefully evaluated over years, especially if virulence to the individual *R* genes occurs in the local pathogen populations. For example, old potato cultivars such as ‘Pentland Dell’ (R1R2R3) and ‘Escort’ (R1R2R3R10) nowadays succumb to late blight since virulent races have evolved [14]. In bread wheat, introduction of a transgene cassette of five resistance gene initially led to resistance to aggressive and highly virulent *Puccinia graminis* f. sp. *tritici* (*Pgt*) isolates, however a new *Pgt* isolate that can overcome several genes from this cassette has already emerged [15]. A similar scenario can be predicted for the *R* gene destroyer, *P. infestans* [3], and its favorite host, potato; it is only dependent on the passage of time.

Unlike ETI, the pattern-triggered immunity (PTI) has not been extensively explored in potato yet. PTI employs the pattern recognition receptors (PRR) to detect pathogen-associated molecular patterns (PAMPs) and trigger immunity [6]. PRRs can be receptor-like kinases (RLKs) or receptor-like proteins (RLPs), depending on the presence or absence of the kinase domain, respectively [16]. PRR-mediated immunity is thought to be more durable, since PAMPs are conserved structures or patterns essential for microbial viability or lifestyle, and shared by different groups of microorganisms [17, 18]. PAMPs are less likely to change and PRR are therefore considered to confer a more stable and broad-spectrum resistance compared to NLR [19, 20]. Resistance activity based on PRR can even be obtained beyond host plant families. The transfer of the PRR EFR from *Arabidopsis thaliana* to *Nicotiana benthamiana* and *Solanum lycopersicum*, based on the recognition of the conserved elf18 peptide, led to increased resistance to diverse plant pathogenic bacteria including *Pseudomonas, Agrobacterium, Xanthomonas* and *Ralstonia* [21].

Recently, we have cloned and characterized a new PRR, called *PERU* (Pep-13 receptor unit) from the *Solanum tuberosum* Group Phureja DM 1-3 516 R44, which is a receptor-like kinase that binds the PAMP Pep-13. Pep-13 is an oligopeptide of 13 amino acids, identified within a cell wall-associated transglutaminase (TGase), that showed to be necessary and sufficient to trigger defense responses in parsley and is conserved among different *Phytophthora* species [22, 23]. In potato, Pep-13 triggers oxidative burst and hypersensitive-like cell death, induces the accumulation of salicylic acid, jasmonic acid and the expression of defense-related genes [24, 25]. PERU interacts physically with Pep-13 and forms a complex with BAK1 upon Pep-13 treatment to trigger defense responses; furthermore, the overexpression and knock-out of *PERU* results in an enhanced resistance and increased susceptibility respectively, which indicates that PERU contributes to the resistance in potato against *P. infestans* [26].

The PRR RLP23 from *Arabidopsis thaliana* recognizes nlp20, a conserved peptide of 20 amino acids in Necrosis and Ethylene-inducing Peptide 1 (NEP1)-like proteins (NLPs) produced by bacteria, fungi, and oomycetes [27, 28]. This peptide is sufficient to trigger immune responses in Arabidopsis and other *Brassicaceae* species [28, 29], and it was shown that ectopic expression of RLP23 in potato enhances immunity to *Phytophthora infestans* [29].

Whereas NLRs typically provide complete, or high levels of resistance to *P. infestans*, PERU and RLP provide lower levels of quantitative resistance [4, 26, 29]. We hypothesize that the combination of various PRRs may generate a more adequate level of resistance. In this study we stacked the newly found receptor PERU and RLP23, and investigated if the presence of two PRRs strengthens the immune responses in potato upon PAMP treatment, and generates a more robust resistance to *P. infestans*.

Potato cultivar Atlantic plants were transformed with pK7WG2::*PERU* and the lines *PERU*#10 and *PERU*#11 were obtained and characterized in a previous study [26]. In addition, cv. Atlantic explants were transformed with pB7WG2::*RLP23*, and transformants were obtained. Potato plants were maintained and clonally propagated *in vitro* on MS medium supplemented with 20% sucrose at 25 °C. Seven transformants were evaluated by PCR using different primer combinations (Table S1), and six events turned out to be positive for the transgene. For phenotyping, leaves of 4-week-old plants were treated with nlp20 and the ROS production was measured. Five lines tested positive, namely *RLP23* #1, #2, #4, #5, and #7 (fig. S1a). Double transformants were generated by co-transformation of cv. Atlantic explants with pK7WG2::*PERU* and pB7WG2::*RLP23*. A total of nine double transformants were obtained and subjected to PCR analysis to verify the presence of both genes. The PCR results confirmed the presence of PERU and RLP23 in all nine lines. Subsequently, six lines were randomly selected for phenotyping. ROS production upon nlp20 (fig. S1b) or Pep-13 (fig. S1c) treatment was measured in these six PCR-positive lines, and 5 of them tested positive for both genes, namely PERU/RLP23 #1, #4, #10, #27, #29 (fig. S1b, c).

To avoid the use of saturating doses, we determined EC_50_ by quantifying the ethylene accumulation using increasing concentrations of the peptides nlp20 and Pep-13. We found that Pep-13 triggers ethylene production in potato plants at nanomolar concentration (EC_50_=2.40 nM) (Fig. 1A), while nlp20 triggers responses at micromolar concentration (EC_50_=1.76 µM) (Fig. 1B). Indeed, the double transformants treated with both peptides showed no increase in ROS production (fig S2a), nor ethylene accumulation (fig. S2b), compared with the responses upon Pep-13 treatment alone, which indicated a saturation of the system. Therefore, based on the EC_50_ results, we decided to use 10 nM Pep-13 and 1µM nlp20 for the following experiments.

**Figure 1.**
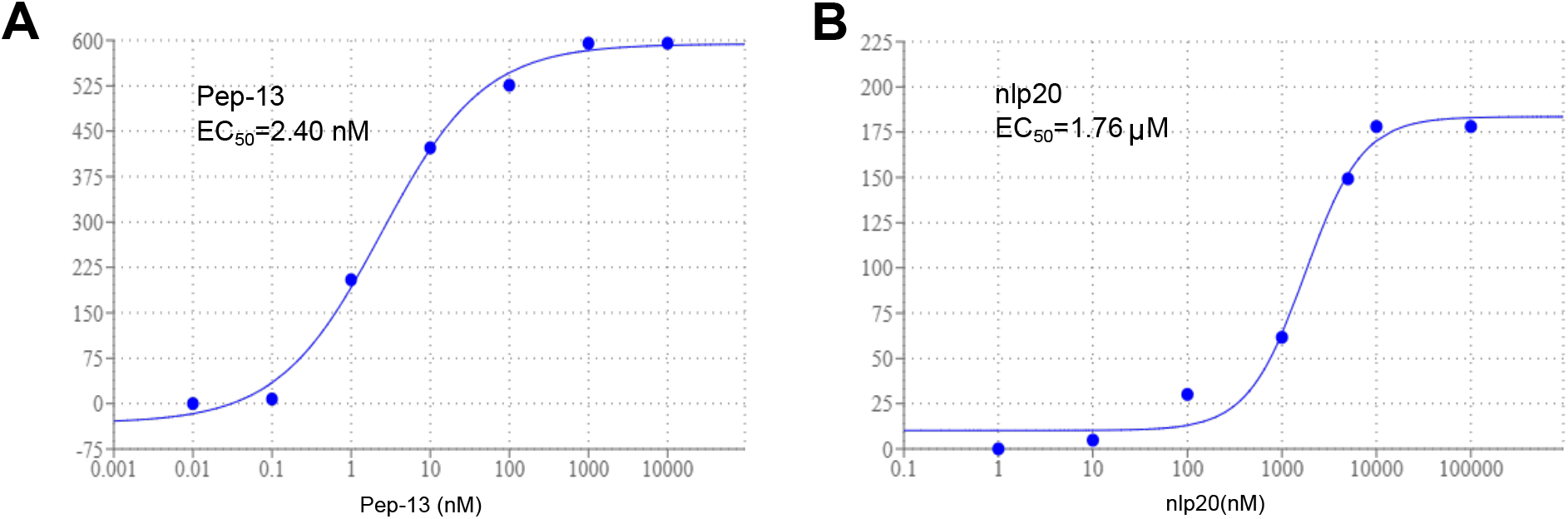
EC_50_ determination of Pep-13 and nlp20 in cv. Atlantic potato plants. Seven different concentrations of Pep-13 (**A**) and nlp20 (**B**) were used to measure the ethylene accumulation in leaves of 4-week-old plants. The plants were much more sensitive to Pep-13 compared to nlp20. The curves and EC_50_ values were obtained in AAT Bioquest.

To test if the presence of two stacked PRRs in potato has an additive effect on the PTI responses to the corresponding molecular patterns, we treated the double transformants with 10 nM Pep-13 and 1 µM nlp20 separately and together, and measured ROS production and ethylene biosynthesis [31]. We observed that in all 4 lines tested, the ROS burst upon Pep-13 treatment is stronger than upon nlp20 treatment, even when Pep-13 is used at nanomolar concentrations (Fig. 2A). This shows that potato cultivar Atlantic is much more sensitive to Pep-13 than nlp20, perhaps because *PERU* can operate in its native background potato, unlike RLP23 that is from Arabidopsis. The treatment of lines expressing *PERU* and *RLP23*, with Pep-13 and nlp20 together, showed a trend of additive effect of PRR stacking on ROS production, which however was not statistically significant (Fig. 2A). We observed this trend in PERU/RLP23 #10, #27, #29, but not in PERU/RLP23 #1. In terms of ethylene accumulation, we found an additive effect upon treatment of combined Pep-13 and nlp20 (Fig. 2B), which was statistically significant for PERU/RLP23 #1, but not significant for PERU/RLP23 #10, #27, #29. Furthermore, we noticed that the treatment of the double transformants with both peptides produced a change in the shape of the ROS burst curve, it showed two phases instead of one, that resembles the junction of curves produced by Pep-13 and nlp20 individually (Fig. 3). To verify that the additive effect is result of the specific recognition of the peptides by their respective receptor, we used the single transformants as controls and treated them with 10 nM Pep-13 and 1 µM nlp20, separately and together. In general, we found that the treatment of single transformants with both peptides, produces no accumulative effect on ROS burst (fig. S3), as expected. Only for PERU #11, we found a minor, not significant increase in ROS when the leaf discs were treated with Pep-13 and nlp20, possibly attributable to variations in the sampled leaves. Besides, the single transformants treated with both Pep-13 and nlp20, showed a normal ROS curve, with only one phase and no changes in shape were observed (fig. S4). Altogether, these results indicate that the stacking of PERU and RLP23 have a small additive but not synergistic effect on the early immune responses like ROS burst and ethylene accumulation.

**Figure 2.**
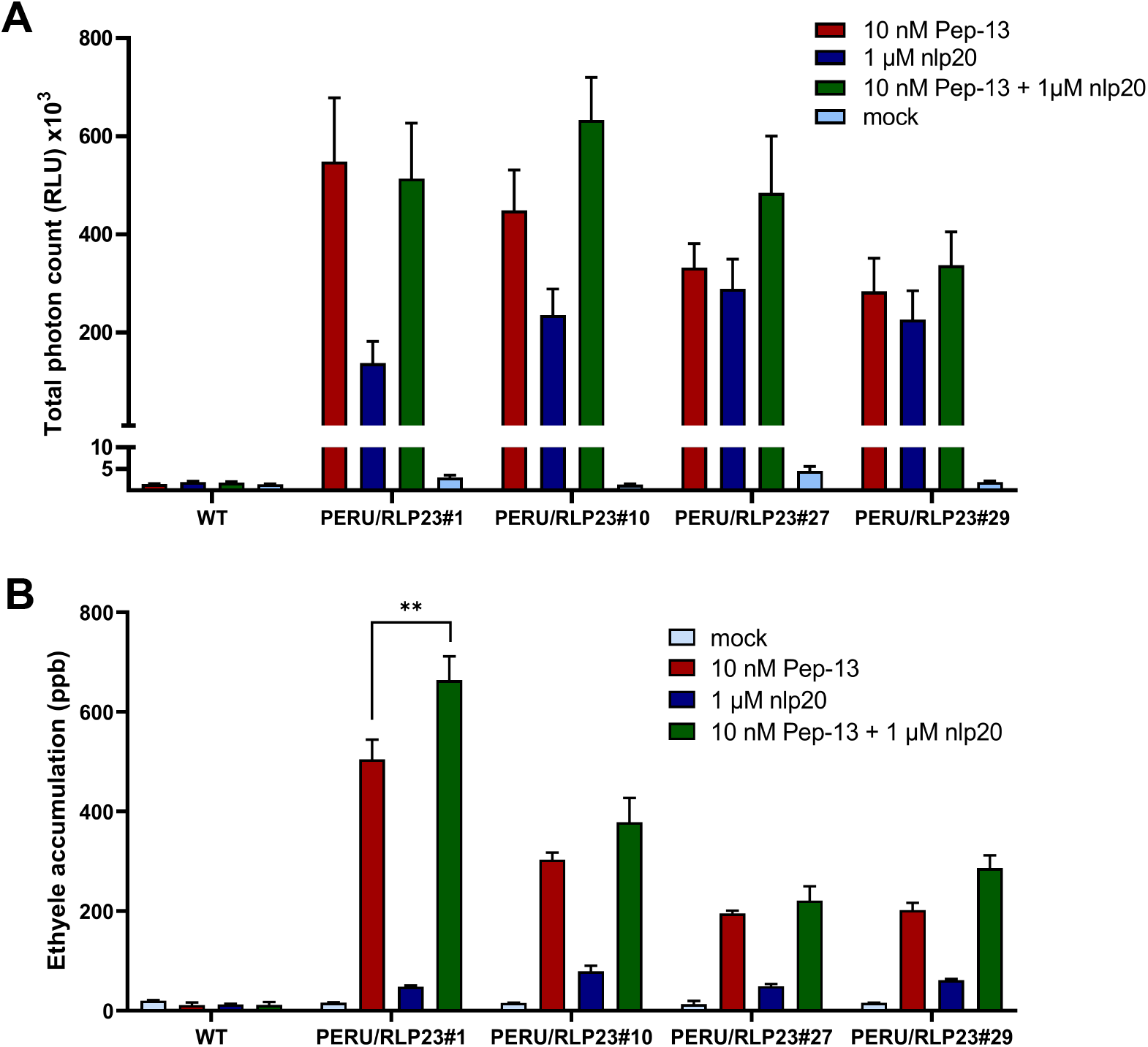
Early immune responses in PERU/RLP23 double transformants treated with Pep-13, nlp20, and Pep-13+nlp20. Leaf discs from four PERU/RLP23 double transformants were selected and treated with 10 nM Pep-13, 1 µM nlp20 and 10 nM Pep-13 + 1 µM nlp20 at the same time. **A**. ROS burst was measured, and the total photon count is shown. An additive effect was observed in the PERU/RLP23 transformants #10, #27 and #29. **B**. The ethylene production was measured, and a significantly higher production was found in PERU/RLP23 #1 in presence of 10 nM Pep-13 and 1µM nlp20 compared to the treatment with single peptides. One-way ANOVA was performed to analyze significant differences between treatments (^**^*P*≤0.01).

**Figure 3.**
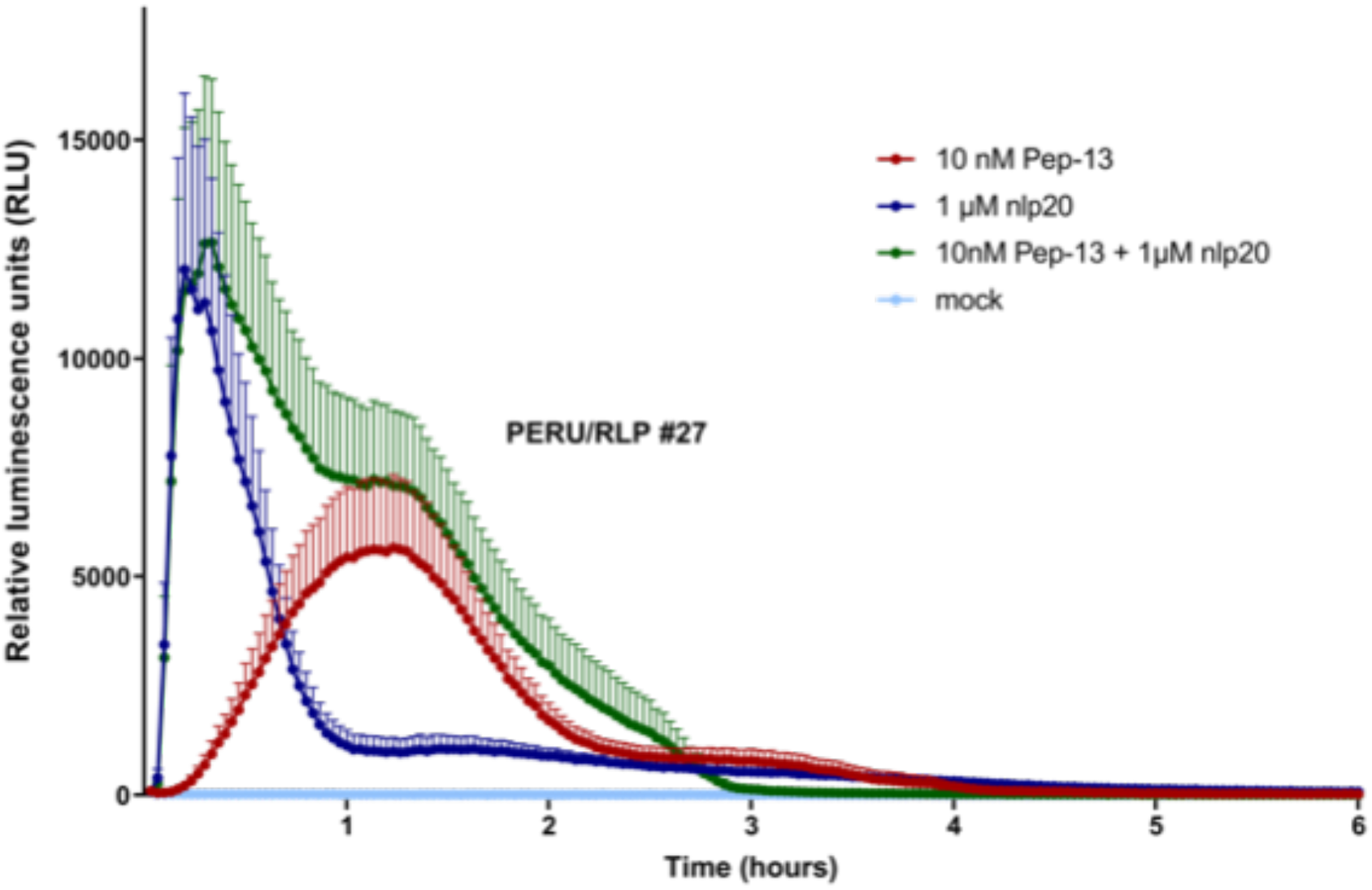
Biphasic ROS curves in PERU/RLP23 double transformant plants. ROS production in transformant PERU/RLP23 #27 treated with Pep-13, nlp20, Pep-13+nlp20, and water (mock). A biphasic curve is produced upon treatment with both Pep-13+nlp20 peptides.

Transformation of individual PRRs has been successfully applied to generate more resistant crops to diseases, although a rather modest level of quantitative resistance is achieved against late blight in potato [21, 29, 32]. We investigated if the stacking of the PRRs *PERU* and *RLP23* in potato can provide a stronger resistance to *P. infestans*. An infection assay was performed as previously described [33], using the WT potato cultivar Atlantic, two single transformant lines per gene (PERU #10, #11, RLP #1, #4), and two double transformant lines (PERU/RLP23 #27, #29). Five plants per genotype, and three compound leaves per plant were spot-inoculated with the highly aggressive *P. infestans* isolate Dinteloord under controlled conditions, and the lesion sizes were measured at 3, 4 and 5 dpi. We observed that *PERU* and *RLP23* contribute to the resistance against *P. infestans*, in line with previously reported findings [26, 29]. The lines overexpressing one PRR or both PRRs are less susceptible to *P. infestans* compared to the WT line, and they showed significant smaller lesion sizes at the three time-points of evaluation (3, 4 and 5 dpi) (P<0.05) (Fig. 4A). The double transformants showed smaller lesions compared to the single transformants at 3dpi, which may indicate that the stacking of the PRRs has a small additive effect on the resistance against *P. infestans* (Fig. 4A). Representative leaves displaying this effect are shown in Fig 4B. At 3 dpi, PERU/RLP23 #29 showed significant smaller lesions, and PERU/RLP23 #27 also showed significant smaller lesions than the single transformants, except for RLP23 #4. At 4dpi, PERU/RLP23 #29 still showed significant differences when compared to the single transformants except for to RLP23 #4, while PERU/RLP23 #27 showed significant smaller lesions compared to PERU #11 and RLP #1. Finally, at 5dpi PERU/RLP23 #29 showed significant smaller lesions compared to PERU #11 and RLP23 #1 only, and PERU/RLP23 #27 showed significant smaller lesions compared to PERU #10,11 and RLP #1. These results suggest a potential role of PRRs stacking to increase resistance to *Phytophthora infestans*.

**Figure 4.**
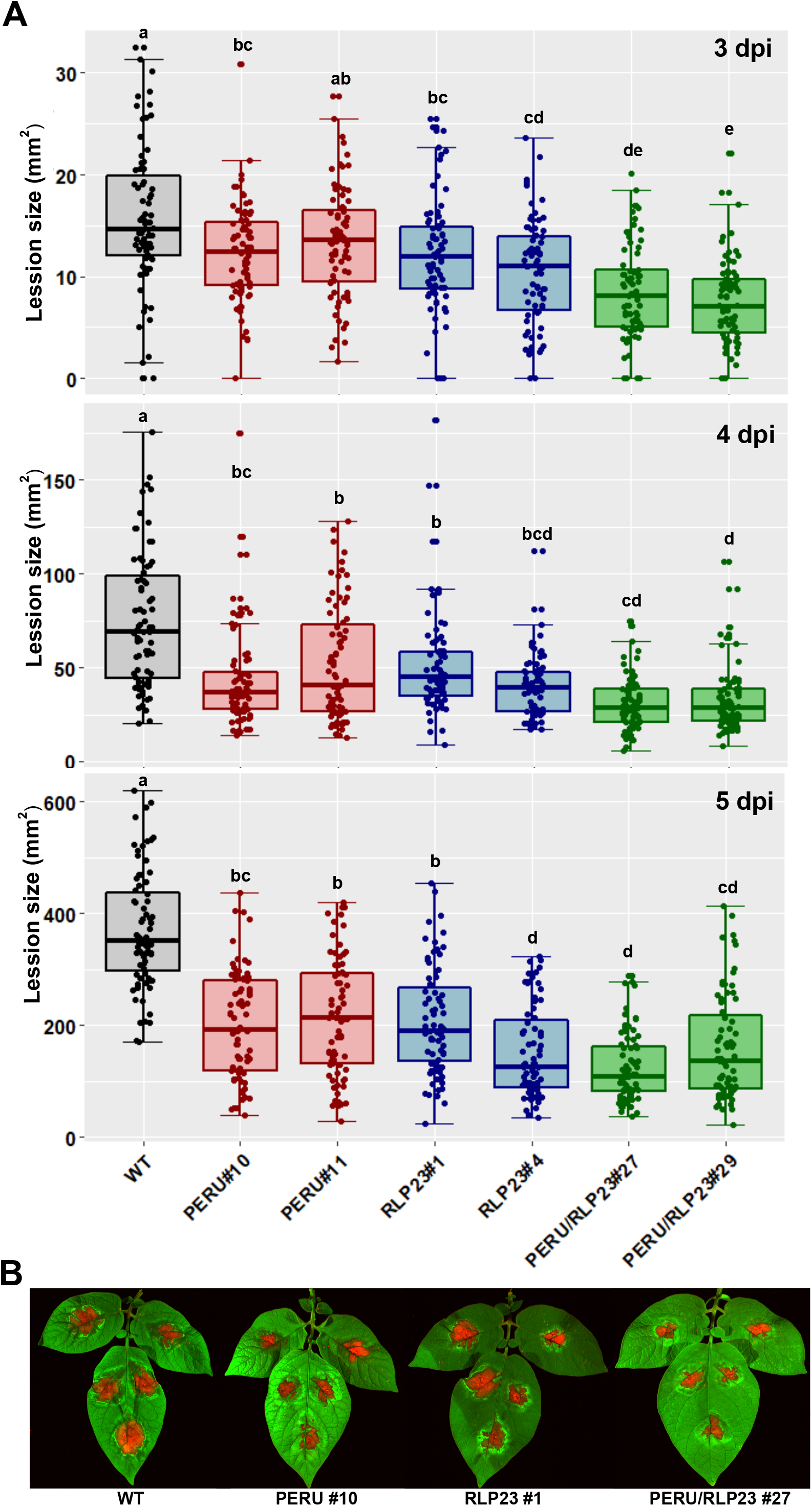
Stacked PRRs have a positive effect on *Phytophthora infestans* resistance. **A**. Leaves of intact potato plants were spot-inoculated with the oomycete *P. infestans*, strain Dinteloord, and lesion sizes (mm^2^) were determined at 3, 4 and 5 dpi **B**. Representative leaves at 4 dpi are shown for Atlantic control (WT), Atlantic-PERU #10, Atlantic-RLP23 #1 and Atlantic PERU/RLP23 #27.

In this study, we hypothesized that the stacking of PRRs could provide more robust resistance to potato against *P. infestans*, an approach that has not been reported so far. We observed that PERU and RLP23 cooperate during early PTI responses like ROS burst and ethylene accumulation upon oomycete patterns treatment (Pep-13 and nlp20), to ultimately slightly enhance the resistance to *P. infestans* infection. Increasing the number of replicates is likely to strengthen the statistical support for our findings.

Additive effects of other PAMPs, elf18 and flg22, on the extracellular alkalinization, were reported previously for Arabidopsis cells, when the peptides were used at non-saturating doses (0.03 nM or 0.06 nM), but not at saturating doses (100 nM or 200 nM) [34]. Additive effect of 1 nM flg22 and 10 nM elf18 on the plasma membrane depolarization was reported as well [35]. In another study, the combined use of different PAMPs showed that certain combinations have additive effect or synergistic effect, while other combinations mutually suppress [36]. The combination of flg22 + elf18, flg22 + LOS (lipo-oligosaccharides from *Xanthomonas campestris* pv. *campestris*), flg22 + core oligosaccharides (also from *X. campestris* pv. *campestris*), and LOS + core oligosaccharides showed a significant increase of calcium ion influx compared to the individual PAMPs, and remarkably flg22 + LOS combination seemed to be synergistic [36]. However, combination of flg22 and oligogalacturonan (OGA) showed a reduced calcium ion influx and ROS burst, compared to the response to flg22 alone. Therefore, for an effective use of PRRs, different combinations of their corresponding PAMPs should be evaluated, to determine if they have an additive, synergistic or antagonistic effect.

Various stacking strategies involving PRRs can be deployed in crops to increase resistance against pathogens. For example, in potato, a PRR and a QTL have been stacked; *EFR* from *A. thaliana* were transferred into a commercial potato line, in which quantitative resistance has been introgressed from *S. commersoni*, showing that *EFR* expression and quantitative resistance have a significant additive effect on the resistance to bacterial wilt caused by *Ralstonia solanacearum* [37]. Secondly, the stacking of PRRs and NLRs, transgenic tomato lines were generated using *Bs2*, an NLR from *Capsicum annuum* for resistance to *Xanthomonas* spp., and *EFR*. As a result, the expression of both genes significantly reduced bacterial incidence and increased the yield [38]. Remarkably, recent studies have shown that PRRs are required by NLR-mediated immunity, NLR-signaling components are required by PRR-mediated immunity, and that these two layers of immunity potentiate mutually [39-42]. Besides this functional relationship, an evolutionary correlation of the number of PRRs and NLRs across plants was revealed [43], which likely implies that PRRs and NLRs are interdependent and that operate synergistically to provide strong immune responses against pathogens. These findings suggests that the transfer of PRRs combined with NLRs is a promising strategy to combat diseases.

Other known PRRs involved in oomycetes recognition could be useful in future experiments, like ELICITIN RECEPTOR (ELR) (from the wild potato species *Solanum microdontum*) or RESPONSE TO ELICITIN (REL) (from *Nicotiana benthamiana*) that mediate the recognition of various elicitins from *Phytophthora* species [32, 44], like RXEG (from *Nicotiana benthamiana*) that recognizes XEG1 of *P. sojae* [45], or RDA2 (from *A. thaliana*) that recognizes 9-methyl sphingoid base from *P. infestans* [46]. Furthermore, several well-known MAMPs and apoplastic effectors from *Phytophthora* remain orphan and their corresponding PRRs are yet unknown, such as CBEL, PcF, SCR74, SCR91, among others [47, 48]. Despite the need of further research, our results are promising and suggest that PRR stacking could be a valuable tool in the battle against *Phytophthora infestans* and other plant pathogens.

## Acknowledgements

This work was supported by the Peruvian Council for science, technology and technological innovation (CONCYTEC) and FONDECYT contract 129-2017 (YCTA). We thank Dirk Jan Huigen for assistance in the greenhouse, Harm Wiegersma and Henk Smid for excellent plant care, Geert Kessel for providing *P. infestans* isolate Dinteloord, and Aviv Andriani for providing advice on disease tests.

