## Supplementary figure 1 for "Stacking of PRRs in potato to achieve enhanced resistance against *Phytophthora infestans*"

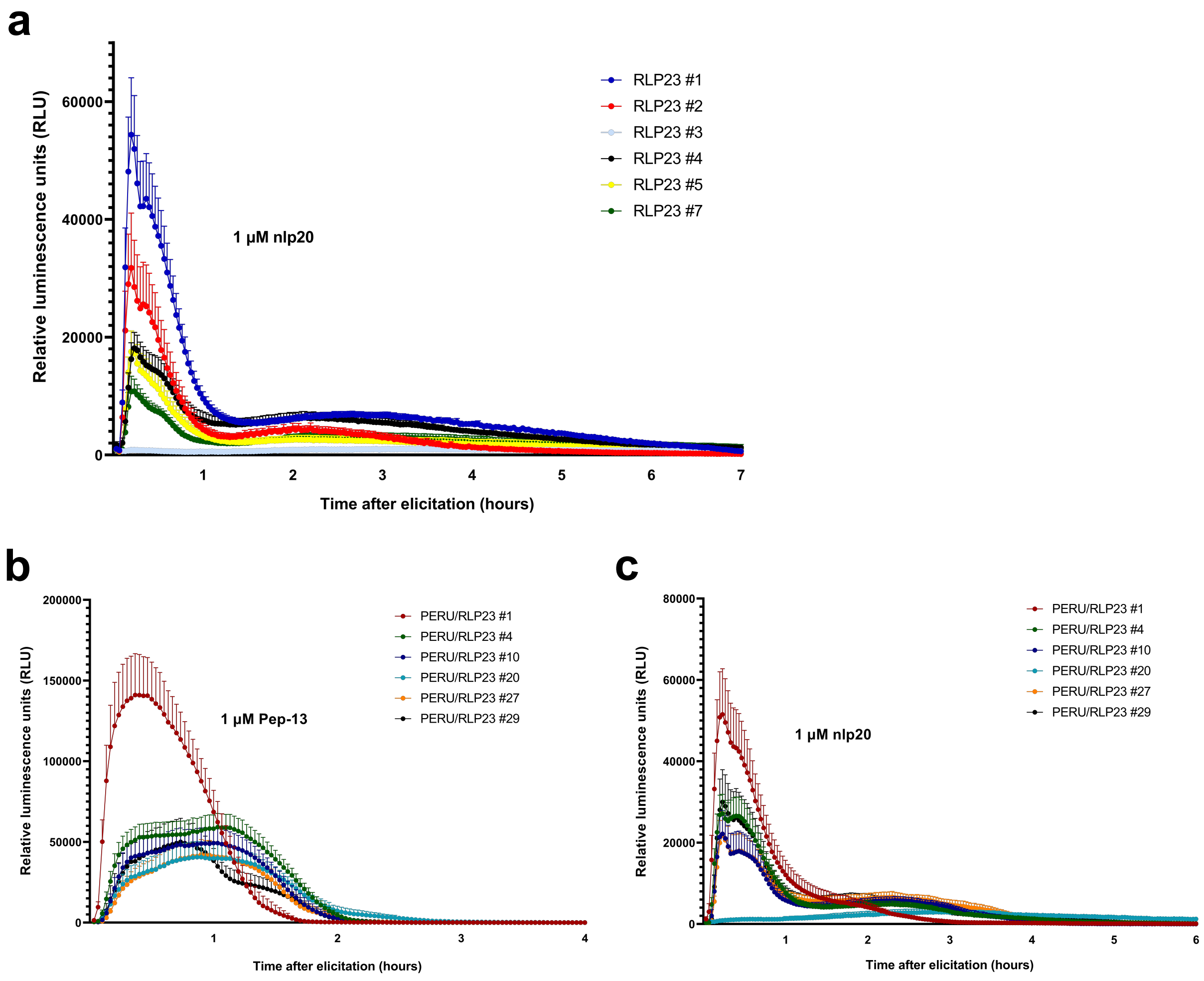


**Supplementary figure 1**. **Stacking of *PERU* and *RLP23* in potato**. **a.** ROS production for single transformants of potato cultivar Atlantic treated with nlp20. Six transgenic lines were treated with 1 μM nlp20, the line RLP23 #3 was genotyped negative by PCR and included here as a control. Five transgenic lines positives for RLP23 presence were identified: RLP23 #1, #2, #4, #5, #7. **b.** ROS production in double transformants of potato cultivar Atlantic treated with Pep-13. Six transgenic lines were treated with 1 μM Pep-13, and all resulted positive. **c.** ROS production for double transformants of potato cultivar Atlantic treated with nlp20. Lines were treated with 1 μM nlp20, all lines except PERU/RLP23 #20 resulted positive.
