## Supplementary figure 2 for "Stacking of PRRs in potato to achieve enhanced resistance against *Phytophthora infestans*"

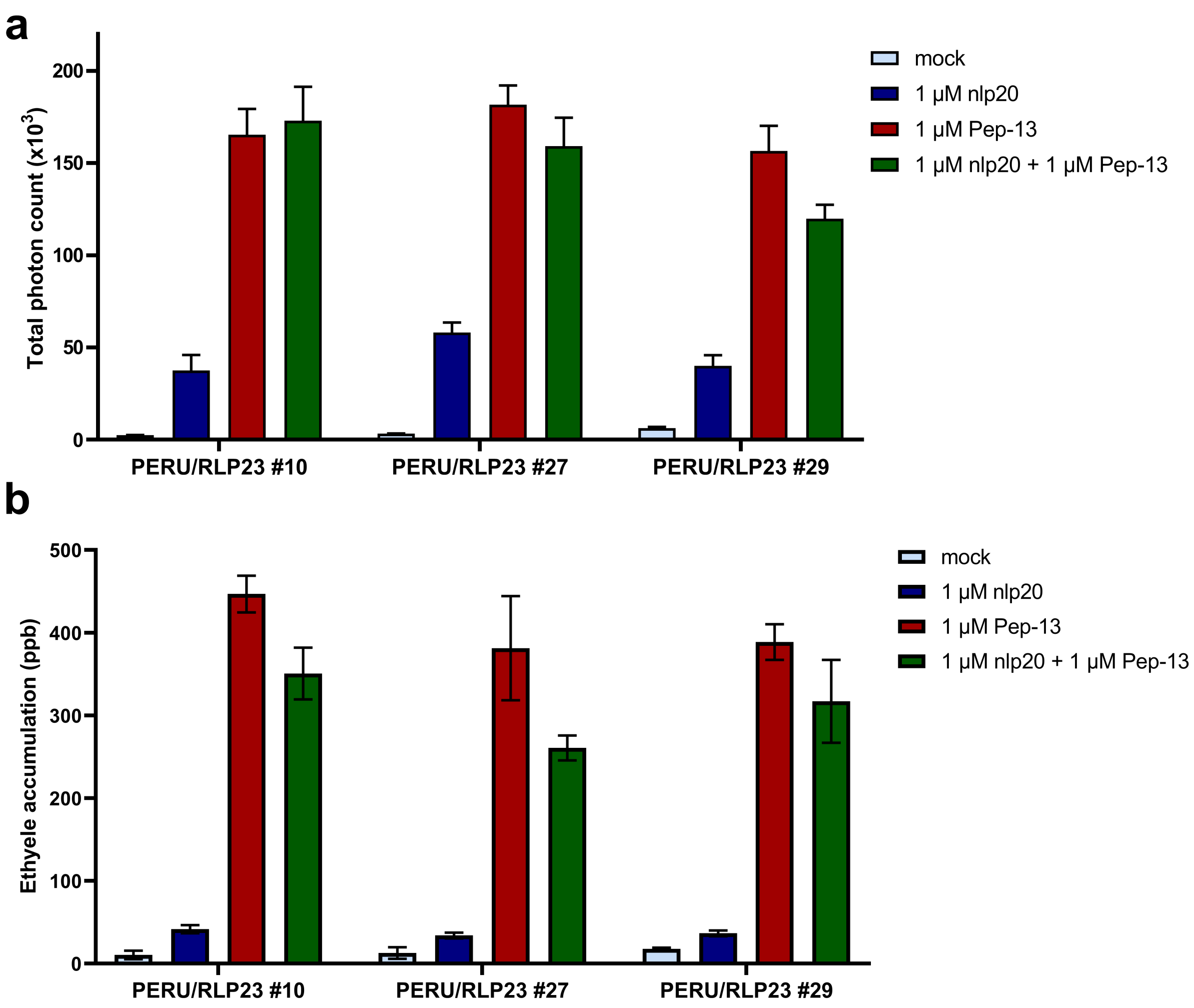


**Supplementary figure 2**. **Early immune responses to saturating doses of PAMPs in PERU/RLP23 double transformants**. **a.** ROS production and **b.** ethylene accumulation in the double transformants PERU/RLP23 #10, #27 and #29 treated with saturating concentration of patterns (1 µM Pep-13, 1 µM nlp20, and a combination of 1 µM Pep-13 and 1 µM nlp20).
