## Supplementary figure 3 for "Stacking of PRRs in potato to achieve enhanced resistance against *Phytophthora infestans*"

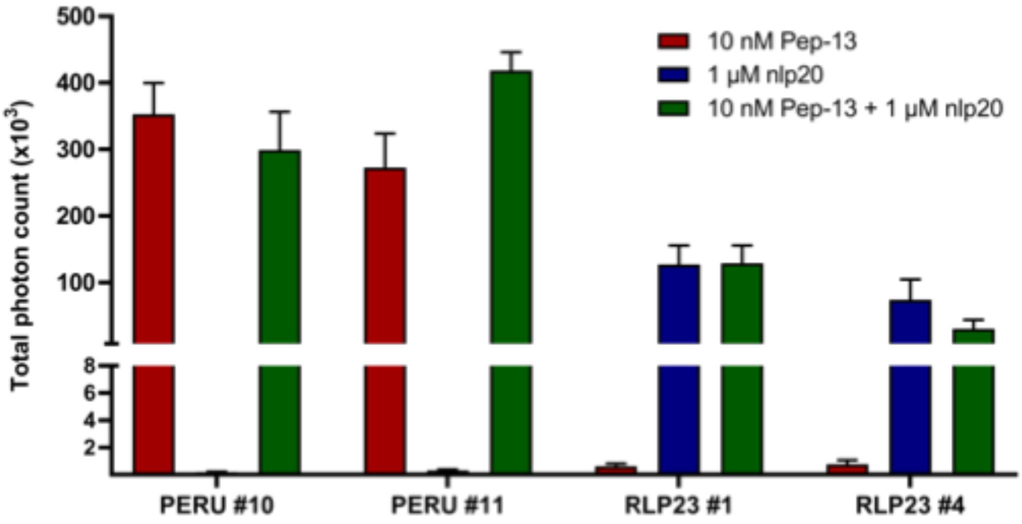


**Supplementary figure 3**. **ROS production in the single transformants.** Total photon count in PERU #10, PERU #11, RLP23 #1 and RLP23 #4 plants treated with 10 nM Pep13, 1 µM nlp20, or a combination of 10 nM Pep-13 and 1 µM nlp20.
