## Supplementary figure 4 for "Stacking of PRRs in potato to achieve enhanced resistance against *Phytophthora infestans*"

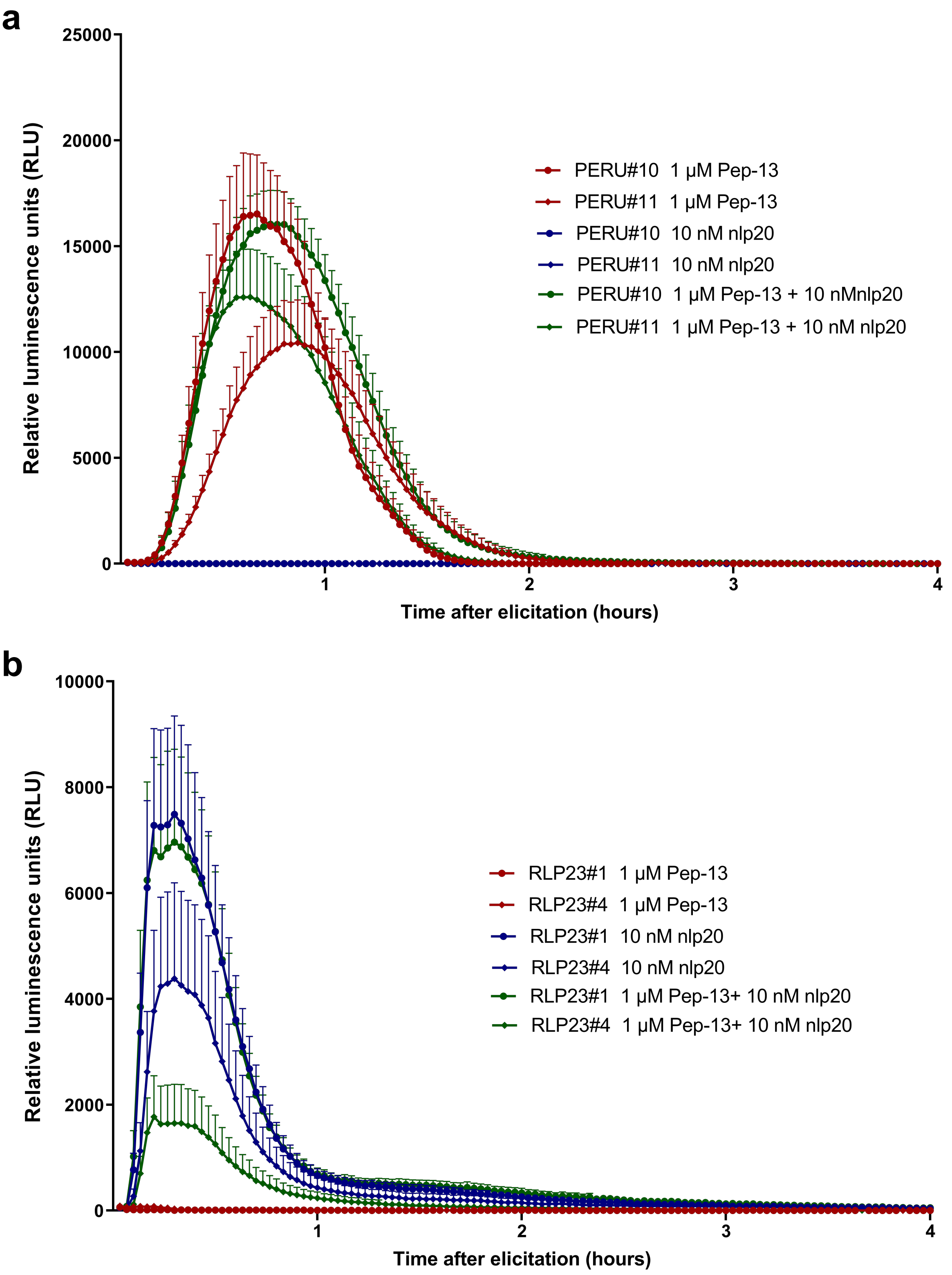


**Supplementary figure 4**. **Curves of ROS burst in the single transformants**. **a.** PERU #10, PERU #11, and **b.** RLP23 #1, RLP23 #4 plants were treated with 10 nM Pep-13, 1 μM nlp20, or a combination of 10 nM Pep-13 and 1 μM nlp20. The measurement of ROS production was conducted over a period of 4 hours.
