## Supplementary Materials and Methods for "Stacking of PRRs in potato to achieve enhanced resistance against *Phytophthora infestans*"

### Supplementary information

### Materials and Methods

**Plant materials.** Potato plants were maintained and clonally propagated in vitro on MS medium supplemented with 20% sucrose at 25 °C. For experiments, 2 weeks old plantlets were transferred to sterilized soil and grown in regulated greenhouse compartments at 18-22 °C, 16h/8h day/night regime and 70% relative humidity.

**Peptides and protein.** Peptides were synthesized by Genscript USA Inc, prepared as 1mM stock solutions in (DMSO), and diluted in MilliQ water or sterile tap water before use. We used nlp20 from *Phytophthora parasitica* (AIMYSWYFPKDSPVTGLGHR) and Pep-13 from *P. sojae* (VWNQPVRGFKVYE).

**Constructs and stable transformation.** *PERU* was cloned in pK7WG2 under the 35S promotor, and *RLP23* was cloned in pB7WG2 [29]. Stable transformation of potato cultivar Atlantic was carried out using routine potato transformation protocols [49]. To generate the single transformants, potato internodes were co-cultivated with *Agrobacterium tumefaciens* AGL1 contained the specific construct. The explants were transferred to a regeneration medium with 100 mg/L kanamycin or 2mg/L phosphinothricin (PPT, glufosinate ammonium) as selection agent. To obtain the double transgenic lines, potato internodes were co-transformed with the two *A. tumefaciens* cultures, and the explants were transferred to a regeneration medium with 100 mg/L kanamycin and 2mg/L PPT. The regenerants were genotyped using specific primers targeting *virG*, selection marker, *35S*, *PERU* and *RLP23*. Subsequently, the plants were phenotyped by ROS to test the response conferred by *PERU* or *RLP23*.

**Measurement of ethylene accumulation**. Leaves of 4-week old potato plants were cut in round pieces (6.0 mm diameter) and floated in MilliQ water overnight. Four leaf pieces were incubated in sealed 10 ml vials containing 0.6 mL 20 mM MES buffer, pH5.6 and the indicated elicitor. Ethylene accumulation was measured after 4 hours of incubation by gas chromatographic analysis (TraceGC1300 (InterScience, Breda, NL) coupled with a flame ionization detector) of 3.5 mL of the air drawn from the closed vial with a syringe. At least, three replicates were measured for each treatment.

**Measurement of ROS production.** Leaves of 4-week-old potato plants were cut in round pieces (6.0 mm diameter), placed in 96 well white plate containing 50 µL of MilliQ water, and incubated overnight. Water was removed and replaced with 50 µL of new MilliQ water. After that, 50 µL of a solution containing 10 µg horseradish peroxidase, 50 µM luminol L-012, and the desired elicitor. Luminescence was measured using a CLARIOstar plate reader (BMG LABTECH) over a period of at least 3 hours.

***Phytophthora infestans* infection assay.** Disease tests were performed as previously described [32]. In brief, *P. infestans* isolate Dinteloord from our in-house collection was grown on rye agar medium supplemented with 20 g/L sucrose at 18 °C in the dark. To obtain zoospores, the mycelium was flooded with cold water (4 °C), the suspension was transferred to a new tube and incubated at 4 °C for approximately 2 hours. The number of zoospores was counted and adjusted to $5x10^4$ zoospores/mL for inoculation. Intact, four-weeks-old plants of potato cv. Atlantic WT along with two single transformants expressing *PERU*, two single transformants expressing *RLP23,* two double transformants expressing *PERU* and *RLP23* were spot-inoculated with zoospore suspensions. Five plants per genotype, and third to fifth fully developed leaves (counted from the top) were used. Three leaflets per compound leaf were inoculated (three spots per main leaflet, and one spot per other small leaflets) by pipetting 10 µl droplets on the abaxial side. Lesion diameters were measured at 3, 4 and 5 dpi. The experiment was performed twice.
