## Supplementary Table 1 for "Stacking of PRRs in potato to achieve enhanced resistance against *Phytophthora infestans*"

**Supplementary table 1**. Sequence of primers used in this study.

| Primer name | Sequence 5' → 3' | Target |
|---|---|---|
| pK7-F | CTATTCTAGTCGACCTGCAG | pK7WG2 vector |
| pK7-R | GAGACTGGTGATTTTTGCGG | pK7WG2 vector |
| 35S-F | TGCTGACCCACAGATGGTTA | CaMV 35S promotor |
| 35S-R | CGGCGAGTTCTGTTAGATCC | CaMV 35S promotor |
| NPTII-F | GCGTTCAAAAGTCGCCTAAG | nptII (kanamycin resistance) |
| NPTII-R | AGTGACAACGTCGAGCACAG | nptII (kanamycin resistance) |
| virG-F1 | GCCGACAGCACCCAGTTCAC | virG virulence gene |
| virG-R1 | CCTGCCGTAAGTTTCACCTCACC | virG virulence gene |
| bar-F | GAGACGTACACGGTCGACTC | Bar gene |
| bar-R | ACGTTGAAGGAGCCACTGAG | Bar gene |
| RLP23-F | CATCCCGTTGACCCTGGAAA | RLP23 gene |
| RLP23-R | AAGAGGCACCATCACAGAGC | RLP23 gene |
| PERU-F | GGCTCTGTCTACAAAGGCGT | PERU gene |
| attR-R | CCGCGGGATATCACCACTTT | PERU gene |
