## Supplementary Table 2 for "Stacking of PRRs in potato to achieve enhanced resistance against *Phytophthora infestans*"

**Supplementary table 2**. Constructs used in this study.

| Construct | Binary vector | *A. tumefasciens* strains | Reference |
|---|---|---|---|
| PERU | pK7WG2 | AGL1 | Torres Ascurra et al. 2023 |
| RLP23 | pB7WG2 | AGL1 | Alberts et al. 2015 |
